# FeatureForest: the power of foundation models, the usability of random forests

**DOI:** 10.1101/2024.12.12.628025

**Authors:** Mehdi Seifi, Damian Dalle Nogare, Juan Battagliotti, Vera Galinova, Ananya Kedige Rao, AI4Life Horizon Europe Programme Consortium, Johan Decelle, Florian Jug, Joran Deschamps

## Abstract

Once the work at the microscope is done, biological discoveries rely heavily on proper downstream analysis. This often amounts to first segmenting the biological objects of interest in the image before performing a quantitative analysis. Deep-learning (DL) is nowadays ubiquitous in such segmentation tasks. However, DL can be cumbersome to apply, as it often requires large amount of manual labeling to produce ground-truth data, and expert knowledge to train the models from scratch. Nonetheless, the performance of large foundation models, although trained on natural images, are improving on scientific images with every new model released. They, however, require either manual prompting or tedious post-processing to selectively segment the biological objects of interest. Classical machine learning algorithms, such as random forest classifiers, on the other hand, are well-established, easy to train, and often yield results of sufficient quality for downstream processing tasks, hence their continued popularity. Unfortunately, they are limited to objects with distinct, well-defined textures compared to their environment. This generally limits their usefulness to structures easy to recognize. Here, we present FeatureForest, an open-source tool that leverages the feature embeddings of large foundation models to train a random forest classifier, thereby providing users with a rapid way of semantically segmenting complex images using only a few labeling strokes. We demonstrate the improvement in performance over a variety of datasets, including large and complex volumetric electron microscopy stacks. Our implementation is available in napari, currently integrates four foundation models, and can easily be extended to any new model once they become available.

## 1 Introduction

Segmentation is an ubiquitous task in microscopy image analysis, as it enables downstream processing and quantification of objects of interest. Researchers have at their disposal a wide array of algorithms, among which machine learning approaches have long been the methods of choice. In particular, random forest pixel classification is a well-established algorithm, at the heart of several popular software tools for bioimage analysis [1, 2, 3, 4]. It uses common image filters to extract a feature vector representation of hand-labeled pixels in order to train decision trees to best match the given input labels. Because the image filters can be 2D or 3D, random forest pixel classifiers can natively perform 3D segmentation. Moreover, they are compatible with multiclass pixel classification. These algorithms owe their popularity to the simple iterative process by which users draw small scribbles to assign a class to a subset of pixels, rapidly train a random forest, and predict results over many images. This swift training procedure allows the correction of mistakes by adding new labels to the training set and training anew. While random forest pixel classification algorithms have a wide application range covering all types of images and modalities, they are limited in their predictive power, and easily confuse different object types that have similar textures [3].

In recent years, deep-learning has emerged as the most powerful approach for image segmentation. Such approaches are most often trained in a supervised fashion, that is to say with a large dataset of manually segmented images as reference. The likes of StarDist [5] or CellPose [6] are go-to tools for image analysts wishing to perform image segmentation. Once trained, these methods often outperform random forest pixel classification [7, 8]. In addition, both methods are compatible with 3D segmentation. Furthermore, CellPose2 [9] introduced user-friendly fine-tuning of models by providing a user interface to correct errors and retrain the selected model, similar to the way random forest classifiers are used. Base models were trained on datasets consisting of various imaging modalities and diverse samples, and are capable of segmenting objects of similar size in a wide range of images. It does not, however, segment multiple classes, and can struggle to effectively segment objects with various shapes and sizes simultaneously.

With more compute power and more data being available, much larger networks are now being trained with astounding results. For instance, Segment Anything Model (SAM) [10], is capable of accurately segmenting biological objects in 2D in both electron and light microscopy images, all the while being trained on a dataset overwhelmingly composed of natural (i.e. every-day) images. To push the boundary of its capabilities, fine-tuning this model with scientific images is being explored [11]. SAM does not natively segment whole images, but rather expects user annotations - also called prompts - in the form of bounding boxes or points as inputs, and returns segmented instances of the annotated objects. While this is a powerful way to enable interactivity, scientific segmentation pipelines preferentially require automated processing of large datasets. Without an accurate and automated way of producing the prompts, SAM applications in bioimage analysis are limited to direct and time-consuming user interactions for each object in the dataset.

Another fruitful research avenue is the use of rich latent spaces as basis for segmentation. Rather than segmenting pixels directly, other approaches, such as MAESTER [12] or DINOv2 [13, 14], train a large network on a different task (e.g. reconstructing masked areas of the image) in order to produce rich feature embedding of the image. These features can then be used to cluster the pixels based on their proximity in this latent space, and identify object classes with these clusters. While enticing, cluster-based features are often limited by the lack of knowledge of how many classes are expected in a given image, and whether these classes cluster meaningfully in the feature space. Moreover, the application of such approaches are so far limited to deep-learning experts due to the complexity of the training process, and success in segmenting scientific images different from those in the training set is not ensured.

In the context of electron microscopy, for instance, certain imaging methods lead to high contrast images with a high density of objects. These datasets are particularly interesting to the community as they potentially enable the observation of unseen biology or the quantification with high resolution of many biological structures. However, segmenting these structures remains an unsolved challenge owing to the complexity and size of the images. De-novo training of supervised deep-learning models is out of reach for many researchers, and few groups have been capable of systematically spending the time and effort to label large volumes [15, 16]. Large deep-learning networks such as SAM are capable of segmenting objects with high accuracy, provided that users generate the correct prompts. Even though SAM requires annotations that are much easier to produce than whole-object masks, the sheer size of the typical electron microscopy volume prevents annotating every object in the dataset. Here, we present FeatureForest, a method that combines the power of large deep-learning models with the simplicity and user-guidance provided by random forest classification algorithms. With FeatureForest, manual labeling can be a matter of minutes, and user-guidance allows segmenting complex objects throughout entire datasets without requiring re-training large deep learning networks. We showcase how FeatureForest fills a gap in the segmentation of large electron microscopy datasets, enabling researchers to segment challenging images. More specifically, FeatureForest uses large foundation models to extract feature vectors corresponding to user-labeled pixels in order to train a random forest algorithm. In this manuscript, we demonstrate the usefulness of FeatureForest over various scientific datasets for which no straightforward or user-friendly algorithm exist, and the improvements it yields over classical random forest classification. We provide an implementation of FeatureForest in an open napari [17] plugin, as well as example scripts and notebooks to perform prediction outside napari (e.g. on clusters).

## 2 Principle

FeatureForest replaces the classical filters of a random forest classifier with large deep-learning models (see Fig. 1a), and extracts the feature vectors used during random forest training from the embeddings that are computed within those networks. The overall iterative training process remains otherwise similar, with users requiring a few iterations of labeling and training before obtaining desired results. FeatureForest currently includes several foundation models: SAM [10], MobileSAM [18], SAM2 [19] and DINOv2 [13] (see Methods for a description of the feature vectors extraction process). Advanced users can extend this list and adapt the model of their choice for use in FeatureForest (see Methods).

**Figure 1.**
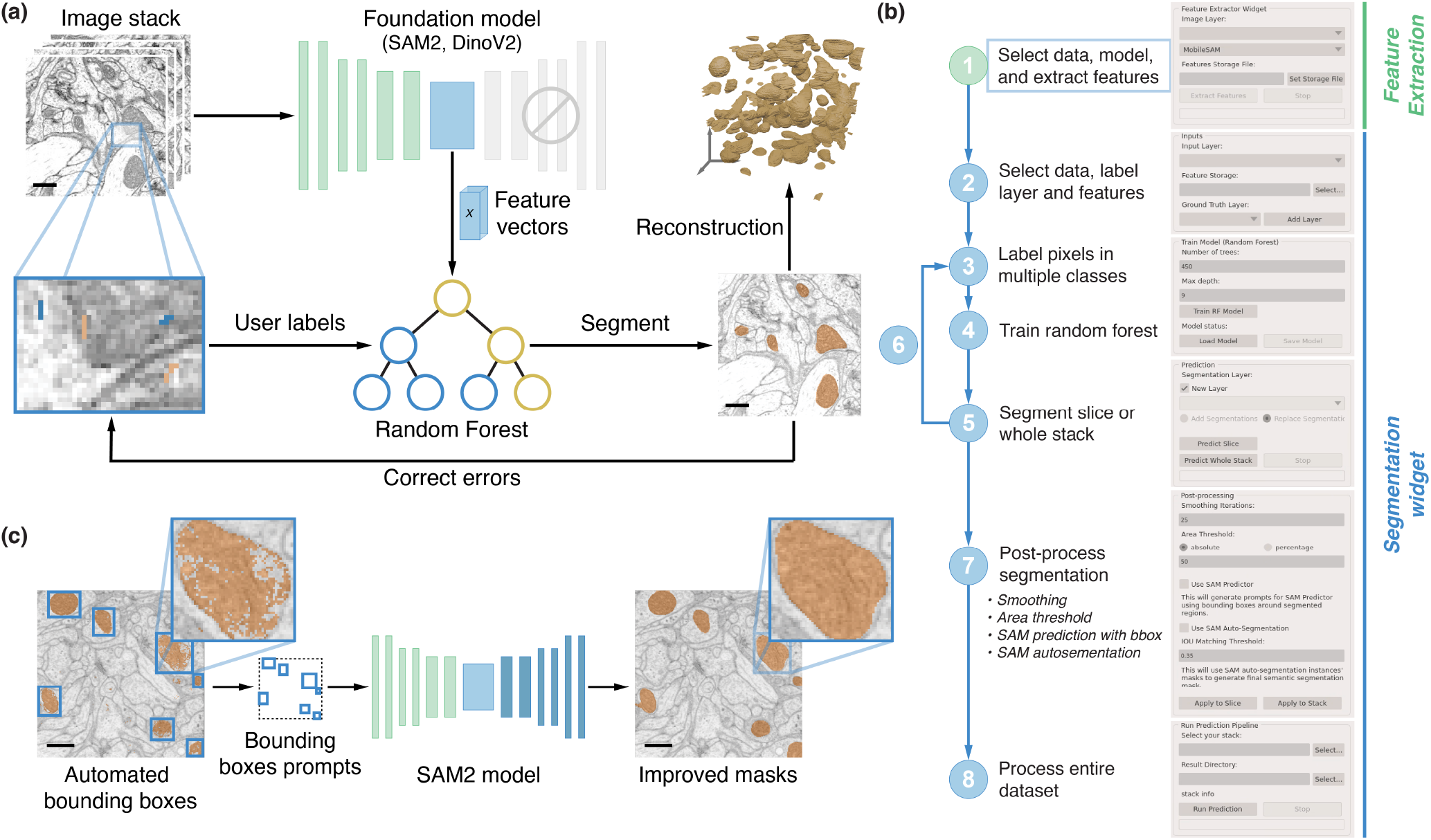
FeatureForest principle. (a) A subset of the data is used to train the random forest model. First, feature vectors from a large deep-learning model corresponding to each pixel in the data are extracted. Users provides both pixel labels and feature vectors to a random forest classifier in order to train the random forest model. Once trained, the classifier can segment slices in the data. An iterative labeling, training and segmenting process is followed to improve the classifier until satisfaction. (b) Overview of the napari widgets and the various steps followed by users. (c) Post-processing with SAM2 is performed by generating bounding boxes around the connected components in the segmentation, and using them as prompts for SAM2. This results in multiple masks that are merged into a semantic segmentation mask.

In Fig. 1b, we describe the FeatureForest pipeline as available to users via the napari plugin we provide. In a first step, using the *Feature Extraction* widget, users extract the feature vectors corresponding to all pixels in a set of images loaded in napari from the model of their choice. The feature vectors corresponding to individual pixels are stored in an HDF5 file to allow random access during the later stages. The feature vectors are large (from 320 to 1536 features per pixel depending on the model) and their extraction slow. For instance, with SAM2 (768 features per pixel), a 512 × 512 slice requires 900 MB of storage space, and required on our system 38 seconds to be generated. Therefore, storing them once for the whole training dataset enables faster iterations when training the random forest. Then, the *Segmentation Widget* is used to train iteratively a random forest on the data subset, as well as perform the final segmentation. First, users select a napari layer containing their data, point to the feature vector file that was exported using the *Feature Extraction* widget, and select their labeling layer. Next, using the napari’s built-in labeling tools, they label a small representative set of pixels before training a random forest on these labeled pixels. Once the training done, users can segment the currently selected slice or full stack. The results can be improved by iteratively adding new labeled pixels where the trained classifier performed poorly. The training process allows rapid iteration between labeling, training and prediction. At any point, users can save the trained random forest classifier. After a few iterations, users can predict on a dataset saved on the disk, e.g. a much larger stack. Furthermore, FeatureForest includes post-processing (see Methods), such as smoothing steps and filtering connected components based on size. Additional post-processing tools leverage SAM2 in two different ways: (i) by generating bounding boxes around instances obtained from performing watershed on the output of FeatureForest and using them as prompts for SAM2 (see Fig. 1c), or (ii) by using the SAM2 auto-segmentation feature in which a grid of points over the image is passed to the model as prompts, and the final masks are selected by thresholding the intersection over union (IoU) between instances obtained from SAM2 and watershed-processed FeatureForest results. Using SAM2 in the post-processing step typically results in object segmentations with smoother boundaries (see Fig. 1c insets).

## 3 Results

### 3.1 FeatureForest on various microscopy modalities

We applied FeatureForest to various datasets from three different imaging modalities: focused ion beam scanning electron microscopy (FIB-SEM), brightfield microscopy, and H&E staining. For each dataset, we trained a classical random forest classifier using Labkit [3] and FeatureForest on the same training images. Additionally, we applied the SAM2 bounding-box post-processing available within FeatureForest (see Methods). In order to quantify the segmentation performance, we computed the Dice coefficient between the resulting segmentation and the ground-truth provided in the public datasets.

FIB-SEM data typically has high contrast and dense structures, while being too large to manually label and too complex to segment using random forest pixel classifiers. Fig. 2a shows a single slice of a fly brain imaged by FIB-SEM as well as the ground-truth masks of mitochondria and the segmentations obtained with Labkit and FeatureForest. The mitochondria appear as dark and round objects of varying intensity. While the random forest classifier is able to classify most pixels from inside the mitochondria, it also creates a high number of false positive and misses their outer membrane. In contrast to the random forest classifier, FeatureForest produces a segmentation with high coverage of the mitochondria and few false positive pixels, which is quantitatively confirmed by a Dice score of 0.56 for the random forest and 0.89 for FeatureForest. Post-processing the segmentation from FeatureForest using the bounding boxes generation and SAM2 (see Fig. 1c) yields smoother segmentation masks and an improved Dice score of 0.93. Similar results are obtained throughout the dataset (see various slices in Supplementary Fig. S1) and computing the Dice score over the entire dataset shows that FeatureForest performs much better than the classical random forest (see Fig. 2b), with mean and standard deviations of 0.87 ± 0.04 (FeatureForest), 0.91 ± 0.03 (FeatureForest + post-processing), and 0.61 ± 0.07 (random forest). In addition to the higher mean Dice score, FeatureForest also results in lower variability and less sensitivity to varying image quality.

**Figure 2.**
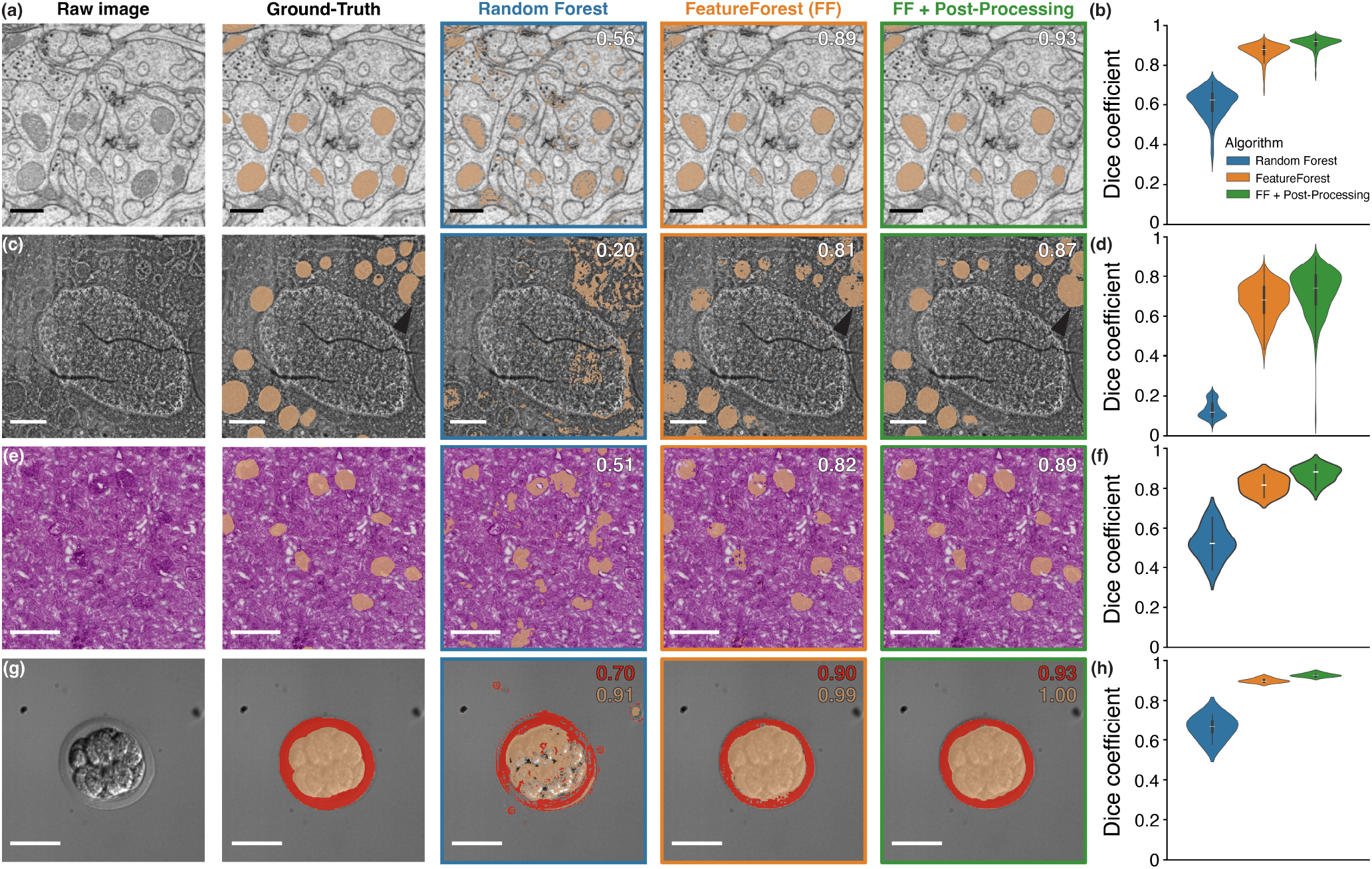
FeatureForest outperforms classical random forest classifiers on complex datasets. (a) FIB-SEM image of a fly brain, overlaid with mitochondria ground-truth mask, and with corresponding segmentation obtained with a random forest classifier, FeatureForest, and after post-processing the results from FeatureForest, from left to right respectively. Dice score with respect to the ground-truth for the specific slice and algorithm is indicated in the top right corner. Scale bar 500 nm. (b) Dice score distribution over the whole dataset (256 slices) presented in (a) for the random forest classifier (blue), FeatureForest (orange) and post-processed FeatureForest (green). (c) FIB-SEM image of a human breast cancer spheroid, overlaid with mitochondria ground-truth mask, and with results from a random forest classifier, FeatureForest, and FeatureForest with post-processing, as in (a). The arrows indicate incomplete segmentation of a mitochondria instance in the ground-truth that is correctly segmented by FeatureForest. Scale bar 1 μm. (d) Distribution of the Dice score corresponding to (c) over the whole dataset (500 slices). (e) H&E staining of a human kidney tissue slice, overlaid with glomerulus ground-truth, and with results from a random forest classifier, FeatureForest, and FeatureForest with post-processing, as in (a). Scale bar 500 μm. (f) Distribution of the Dice score corresponding to (e) over the whole stack (12 tiles). (g) Brightfield image of a mouse embryo, overlaid with cells (orange) and extraembryonic membrane (red) ground-truth masks, and with results from a random forest classifier, FeatureForest, and FeatureForest with post-processing, as in (a). Dice score is indicated for each class. Scale bar 50 μm. (h) Distribution of the Dice score corresponding to the chorion class (red) in (g) over the whole stack (5 slices).

The example dataset of Fig. 2a is a relatively easy segmentation challenge as the mitochondrial texture is sufficiently different from the rest of the image to be well captured by classical image filters. Classical image analysis can further improve the segmentation obtained with the random forest classifier, for instance by filtering connected components by size and applying smoothing or morphological operations. In Fig. 2c, we use another FIB-SEM dataset (human breast cancer spheroid) in which the mitochondria have similar texture to their surrounding and can only be segmented by considering their larger context and shape. Such a situation is exactly where random forest classifiers typically fail, and indeed the classical approach applied to this dataset resulted in a poor quality segmentation (Dice score of 0.20). In comparison, FeatureForest leads to the correct segmentation of the mitochondria with few spurious segmented pixels (Dice score 0.81). As before, the results can be further improved by using our post-processing (Dice score 0.87). The distribution of Dice scores over the whole dataset (500 slices) further shows that FeatureForest enables segmenting the stack with high fidelity while the random forest classifier leads to poor quality results (see Fig. 2d), with mean and standard deviations of 0.67±0.09 (FeatureForest), 0.72 ± 0.11 (FeatureForest + post-processing), and 0.13 ± 0.04 (random forest classifier). The Dice score for FeatureForest decreases throughout the stack (see Supplementary Fig. S2), mostly due to false positive segmentation within the nucleus, which could easily be removed by independently segmenting the nucleus itself and subtracting it from the mitochondria segmentation.

Next, we compared segmentation performance on data from a different imaging modality and sample type. Fig. 2e showcases the output of Labkit and FeatureForest on an H&E stained human kidney tissue. This data contains specific blood vessel structures called glomeruli. In the example from Fig. 2e, the glomeruli are slightly darker than their surrounding and, most importantly, display a wide variety of textures. The random forest classifier is capable of approximately segmenting many glomerulus instances, but misses several of them and produces many spurious groups of segmented pixels (Dice score 0.51). Here again, FeatureForest correctly segments all structures, and its post-processing leads to smooth and complete segmented objects. The dataset was created by tiling a larger image, and some tiles are shown in Supplementary Fig. S3, including the recomposed image, showcasing the performance of FeatureForest. Computing the Dice score for each tile (Fig. 2f) leads to mean and standard deviations of 0.81± 0.04 (FeatureForest), 0.87 ± 0.04 (FeatureForest + post-processing), and 0.52 ± 0.08 (random forest).

Because FeatureForest uses a random forest as classifier on top of the foundational model features, FeatureForest can segment multiple classes at a time. To demonstrate this, in Fig. 2g, we segment a mouse embryo imaged in brightfield microscopy. While the cells at the center of the embryo have a vastly different texture from the rest of the image, the extraembryonic membrane of the embryo has spatially varying intensity due to shadowing and is closer to the uniform background texture. The random forest classifier performs well on the cell mass (Dice score 0.91), but is subpar on the extraembryonic membrane (0.70), leading to incomplete segmentation of the latter. Once again, the classical random forest approach also erroneously segments other structures in the image. FeatureForest produces an almost perfect segmentation with high Dice scores (0.99 for the cells, and 0.90 for the extraembryonic membrane). This is the case throughout all the test images (see Fig. 2h), with mean and standard deviations of 0.90 ± 0.01 (FeatureForest), 0.93 ± 0.01 (FeatureForest + post-processing), and 0.66 ± 0.05 (random forest classifier). To further showcase multiclass segmentation, we also segmented a FIB-SEM dataset distinguishing 6 classes (endoplasmic reticulum, golgi, mitochondria, lysosomes, lipid droplets and nuclear envelope). FeatureForest correctly segments most objects in the images (see Supplementary Fig. S4), across a wide range of texture and shapes.

### 3.2 FeatureForest enables biological discoveries

As we have seen, for complex datasets, the performance of classical random forest pixel classification can lead to unusable segmentation, such as the one shown in Fig. 2c. When training deep learning networks is not possible due the ground-truth label generation requirement, FeatureForest provides a useful alternative to perform the segmentation.

This was exemplified in a recent study [20], in which FeatureForest was used to segment organelles in a complex symbiotic interaction between eukaryotic cells. The data consisted of large resin-embedded FIB-SEM stacks representing a dinoflagellate cell (referred to as the host). This dinoflagellate species is known to acquire and hijack organelles from its algal prey (microalga *Phaeocystis antarctica*), including nucleus, plastids and mitochondria, and retain them over several months.

In Fig. 3a, we compare the manual segmentation of three classes (algal plastids, algal mitochondria and host mitochondria) with the results from FeatureForest on three different slices of a single FIB-SEM stack from [20] (original stack of size 3598 × 4455 × 3944 pixels, which was binned with a factor 4). The mitochondria of both the host (orange) and the algal prey (red) were segmented in two different classes in one FeatureForest model, while we trained FeatureForest again separately for the algal plastids (blue). In all cases, FeatureForest led to high quality segmentation. In particular, the plastids are accurately segmented throughout the stack. To quantify this, we manually segmented 7 test slices distributed over the whole range of the stack. We then computed the Dice score between the manual segmentation and FeatureForest + post-processing on these test slices, confirming the visual impression, with mean and standard deviations of 0.58 ± 0.06 (algal prey mitochondria), 0.64 ± 0.03 (host mitochondria), and 0.88 ± 0.02 (algal plastids) (see Supplementary Fig. S5 for the distributions). Here, manually annotating 7 slices for quantification purposes was a slow process. In contrast, the trained FeatureForest classifier does not require additional inputs to segment the three classes in the 3598 slices of the entire stack. Segmentation of these organelles throughout such a large stack is essential to visualize and quantify morphological changes (e.g. changes in volume and surface of stolen organelles). The segmentation provided by FeatureForest allows building a 3D model of the distribution of organelles in space (see Fig. 3c), a necessary step in measuring the morphometrics of the various organelles. More details on the findings of the study are available in Rao *et al* [20].

**Figure 3.**
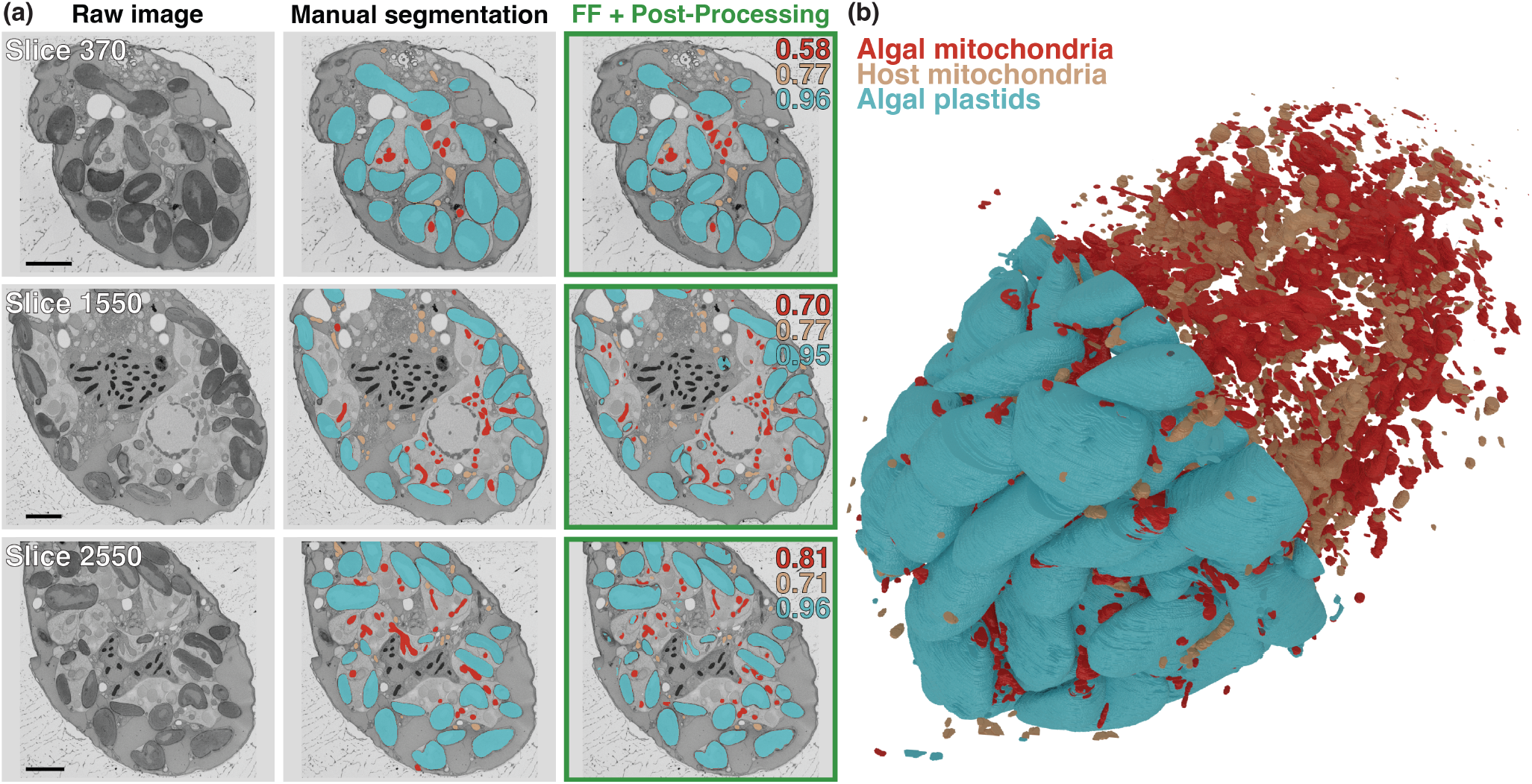
Segmentation of plankton organelles from a FIB-SEM stack using Featureforest. (a) Three different slices (out of 3598) of a dinoflagellate cell imaged in FIB-SEM, overlaid with manual segmentation, and post-processed FeatureForest. The segmentation masks consist of three classes: algal plastids (blue), algal mitochondria (red) and host mitochondria (orange). Dice score between the ground truth and FeatureForest + post-processing is indicated on the top right corner for each class. Scale bar 4 μm. (b) 3D reconstruction of the three classes - algal plastids, algal mitochondria, and host mitochondria - of (a) throughout the entire dataset.

## 4 Discussion

In this manuscript, we introduce FeatureForest, an approach leveraging foundation models to generate high-quality feature representations of pixels which are then used to train a random forest. Via our napari plugin implementation, FeatureForest provides a simple, intuitive and straightforward segmentation pipeline, combining the power of large deep learning image segmentation models with the ease of use of random forests. Crucially, these models can be applied even by researchers with no knowledge of deep learning. FeatureForest fills a gap in the landscape of segmentation tools, in particular for large and complex datasets such as electron microscopy volumes, for which the annotation effort required to assemble ground-truth for deep-learning is considerable. We provide several different foundation models for feature generation, including SAM2, the current state of the art large foundation model for segmentation, as well as the possibility for advanced users to add their own model adapter to FeatureForest. Moreoever, we also designed post-processing steps allowing further improvement in the results of FeatureForest by using its segmentation output to directly generate specific SAM2 predictions.

We benchmarked FeatureForest on multiple publicly available datasets that were published with ground-truth (or for which we could generate our own ground truth), including FIB-SEM, H&E stainings and brightfield images, both for single and multi-class segmentation. We showed that not only does FeatureForest significantly improve segmentation performance on these datasets compared to a classical random forest pixel classifier, but that it also produces high quality segmentation for complex datasets for which the random forest classifier pixel results are unusable.

In our experiments, FeatureForest post-processing improved the results by leading to smoother masks. Overall, the output of SAM and SAM2 tend to follow the same image features than the segmentation obtained by FeatureForest, albeit with more complete connected components. In certain cases, post-processing with bounding box generation can lead to oversegmentation when object instances are difficult to separate using watershed in the FeatureForest segmentation result, and even in rare cases to a mask covering the entire image. In such cases, users might need to post-process these images separately with different parameters (e.g. smaller or higher number of smoothing steps), or implement their own post-processing.

Our method has several limitations that are inherent to the large deep-learning models we are using to extract feature vectors. Firstly, SAM2, SAM and DINOv2 are trained on natural images (e.g. scenes of everyday life, often RGB images) and the feature vectors they produce might not be optimized to separate the biological objects of interest. To address this, fine-tuning these models on microscopy images is an exciting perspective [11].

A further limitation concerns the storage and generation of feature vectors. In order to be time-efficient, feature extraction should preferentially be performed on a graphical processing unit (GPU). Without access to a GPU, users should expect the feature extraction, and the segmentation of full stacks for which features were not pre-exported, to take from minutes to hours depending on the stack size. As this is the most time consuming step, we separated the feature extraction and training steps in the napari plugins. Once FeatureForest is trained, the features are computed on the fly while segmenting an entire dataset. We therefore advise users to train on a representative substack of the image in order to minimize the footprint on disk and generation time of the feature vectors, and segment on the larger stack once they are satisfied with the results on the training stack. In addition, the more complex models have a larger memory footprint as they consist of a much larger number of parameters. For users with limited GPU memory, we also provide lighter models (e.g. MobileSAM [18]) that nonetheless perform well. Future updates will include further optimization of memory usage.

During the development of FeatureForest, similar approaches have been demonstrated, highlighting the usefulness of the method [21, 22, 23, 24]. Compared to these variants, we use state of the art foundation models to generate the feature vectors (e.g. SAM, SAM2), rather than simpler and older networks such as VGG16 [25] or custom networks. Furthermore, we showcase FeatureForest on large and complex datasets, with focus on microscopy imaging, for which available tools for segmentation are limited.

In the future, we will continue to optimize FeatureForest in order to improve user experience, in particular with respect to speed and memory efficiency, and add newer models for feature extraction or post-processing. The source code for the napari plugin is freely and openly available on Github [26], and can be installed through PyPi. We also provide documentation on how to use FeatureForest, as well as scripts and notebooks examples for running FeatureForest outside napari (e.g. on high performance computing (HPC) systems). We believe that FeatureForest constitutes a promising tool for many studies that deal with complex images and for which either knowledge or time prevents researchers from training their own deep-learning segmentation algorithms.

## Methods

### 4.1 FeatureForest

FeatureForest is a Python software package and consists of convenience functions and a napari plugin. All code and documentation is accessible on Github (juglab/featureforest). The FeatureForest napari plugin contains two different widgets: *Feature Extraction* and *Segmentation widget*. The first plugin extracts feature vectors for each pixels in a selected napari layer and stores them in a HDF5 container to allow random access. The second widget allows training the random forest classifier using the previously exported feature vectors, as well as perform post-processing and segmentation of the entire dataset.

#### 4.1.1 Models

The embeddings of deep-learning networks are often of different dimension that those of the input images. In order to obtain per image pixel features, we use patches of smaller size than the expected input size to the network and scale them up. Finally, the resulting feature vectors have smaller spatial dimensions than the original patches and are themselves also up-scaled. This forces the features to have better spatial resolution than without patching.

FeatureForest includes the following models: *SAM2_Large* [19], *SAM2_Base* [19], *SAM* [10], *MobileSAM* [18], and *DINOv2* [13]. All models are implemented by extending the *BaseModelAdapter* class, which allows setting patch size compatible with the specific model, as well as extracting feature vectors for each pixel provided to the model. Each model has its own implementation, as they have different input requirements and architectures.

More specifically, *SAM2_Large* uses “*sam2.1_hiera_large.pt*” as weights, while *SAM2_Base* corresponds to the lighter “*sam2.1_hiera_base_plus.pt*” (see facebookresearch/sam2 on Github). To extract SAM2 embeddings, the images are patched in overlapping patches of size equal to a power of two, and an overlap of half the patch size. The patches are then scaled to 1024*x*1024 before being used as inputs to SAM2. The feature vectors are computed as the concatenation of the image encoder output, leading to 768 features per pixel. The image encoder output has a spatial component, and the tensors are cropped to the non-overlapping regions and scaled back to their original size. *SAM* model uses “*sam_vit_h_4b8939.pth*”. The model differs from SAM2 by the number of output features (1536 per pixel), which are concatenated from both the encoder output and the patch embedding layer. For both SAM and SAM2, the RGB input is simply a concatenation of the same gray-level microscopy image input.

*MobileSAM* model uses a modified version of the *TinyVIT* model architecture that give access to the internal embeddings computed by the encoder. We use “*mobile_sam.pt*” (see ChaoningZhang/MobileSAM on Github) as weights to our modified visual transformer architecture. *MobileSAM* leads to the 320 features per pixel as *SAM*.

Finally, we use “*dinov2_vits14_reg*” from the PyTorch Hub for *DINOv2*. DINOv2 input patches of size divisible by 14. To obtain per pixel output, we create patches of size 70*x*70 with overlaps 28*x*28. The number of output features for each pixel is 384, and is the output of the model itself.

#### 4.1.2 Training

FeatureForest trains a random forest classifier using the feature vectors extracted from one of its adapted models. For each labeled pixel in the labeling layer in napari, the corresponding feature vectors are extracted, and fed along with the label number to the random forest classifier [27]. By default, we use 450 trees of maximum depth 9. The trained classifier can then be used to predict pixel label class for each pixels in the image or slice currently displayed in napari, or predict on the whole stack.

#### 4.1.3 Post-processing

As part of FeatureForest pipeline, we provide several post-processing options that leverage the large deep learning network used for feature generation. In any case, the first step employs mean curvature smoothing, an iterative edge-preserving smoothing method that fill small holes, and filters out small connected components. Users can change the number of smoothing iterations and the threshold used to filter out connected components by area (absolute or relative). By default, we use 25 smoothing iterations, and an absolute threshold of 50 pixels.

Subsequently, users can use either of two additional steps: *SAM2ImagePredictor* and *SAM2AutomaticMaskGenerator*. In the former, we use a watershed algorithm to separate the mask into instances. Bounding boxes are then generated around each instance, and used as prompts for SAM2. The output instances are merged into a single mask and added into napari as a layer. *SamAutomaticMaskGenerator* generates a evenly-spaced grid of points as prompts to SAM2, which outputs a large number of masks. We retain only instances with an intersection over union with respect to the closest connected component from the random forest segmentation larger than a user-set threshold (by default 0.35).

### 4.2 Data and analysis

For each experiment, FeatureForest was run from the commit *97d880d*, with the codebase being available on Github (juglab/featureforest). Unless otherwise indicated, the training and post-processing were carried out with defaults parameters. Labkit [3] was used as the random forest classifier. All training, analysis, and plotting were performed in Python, using the GPU conda environment provided in the source code repository, on a Linux virtual machine with access to a NVIDIA A40-16Q (16 GB) GPU.

The fly brain stack (Fig. 2a and supplementary Fig.S1) is available as part of the *EMPIAR-10982* dataset, and consist of a stack of size 256 × 255 × 255 and an isotropic pixel size 12 nm. We use every 16 frames, starting from the first one, as training set, while prediction was performed on the whole dataset. In the figures, only images that were not used for training and are as far as possible from neighboring training slices are shown. In Fig. 2a, slice number 72 is shown. Dice coefficients in Fig. 2b are computed over the whole stack.

The human breast cancer spheroid stack (Fig. 2c and supplementary Fig.S2) is extracted from *EMPIAR-11380* (sample *F059_bin2*) [28]. The stack has dimensions 1446 × 1683 × 1928 and an isotropic pixel size of 20 nm. We cropped the data to size 500 × 512 × 1024 from the top-left coordinate (390, 800, 150). Training was performed using every 30th frame, starting from the first, while prediction was performed on the whole dataset. In the figures, only images that were not used for training are shown, selecting specifically slices that are as far as possible in z from the training slices. For Fig. 2c, we cropped the slice (slice number 435) to a square region. Dice coefficients in Fig. 2d are computed over the whole stack.

The human kidney tissue example (Fig. 2e and supplementary Fig. S3) is part of a dataset that was compiled from the Human Biomolecular Atlas Program (HuBMAP) and publicly released as part of a Kaggle challenge (https://www.kaggle.com/c/hubmap-kidney-segmentation/data). Specifically, we selected the *1e2425f28* sample, and used the fourth series (resolution 4027 × 3347), and cropped it to 1024 × 3072 (top-left coordinates (486, 1532)), before tiling it into a set of 512 × 512 images (*N* = 12). The masks were provided as instances in a json file and were converted into a binary image, before being cropped and tiled as the raw image. We trained FeatureForest on the first four frames, and predicted on the whole tile stack. We only show in the figures tiles from the range 5-12 (Fig. 2e and supplementary Fig. S3a), and the whole crop in supplementary Fig. S3b. Dice coefficients in Fig. 2f are computed over the whole tile stack.

The mouse embryo dataset (Fig. 2g) is publicly available on the Broad Bioimage Benchmark Collection with access number *BBBC003*. It consists of 5 slices of a 3D brightfield stack of size 640 × 480 and pixel size 420 nm. As the initial ground truth only included the segmentation of the embryo as a single class, we manually labeled the extraembryonic membrane as a second class to generate two-label ground truth. Training was performed on the first slice, and prediction on the whole stack. The fourth slice is shown in Fig. 2g, and is cropped to a square image. Dice coefficients in Fig. 2h are computed over the whole stack.

The U2OS FIB-SEM dataset (Fig. S4) is publicly available as *EMPIAR-11746* [29], and consists of a 1168 × 3394 × 1385 stack with pixel size 2.5 nm in X and Y, and 0.5 nm in Z. We down-scaled the whole stack to a width of 1200, and used every 40 images from slice 500 as training dataset, and predicted on every 30 slice from slice 501 (test dataset). We used 6 out of the 8 classes available in the dataset ground-truth. Slices in Fig. S4 were slightly cropped to exclude white border without information.

The dinoflagellate FIB-SEM dataset (Fig. 3) is part of a recent publication [20]. It was high-pressure frozen and resin-embedded before imaging, and has dimensions 3598 × 4455 × 3944 pixels. More details about sample preparation are available in Rao *et al*. We binned the stack with a factor 4 (3598 × 1113 × 986 pixels) to work on a smaller stack. To obtain the segmentation of the three classes (host mitochondria, algal mitochondria and algal plastids), we trained two different FeatureForest classifiers: one to segment the two types of mitochondria, and one to segment the plastids. In both cases, we use slices 50, 275, 462, 752, 1024, 1375, 1721, 2015, 2310, 2813, and 3067 for training and predicted on every 3 slices (total number of 1200 slices). To allow for quantification, we manually labeled 7 slices (370, 650, 900, 1550, 1850, 2175, 2550) with the three classes using the SAMJ ImageJ plugin. The Dice scores were computed over this test stack. In order to visualize the segmentation, we performed segmentation and post-processing of the three organelles using the classifiers trained for Fig. 3a on the 1200 prediction slices. The final post-processed stack was curated to re-segment with different post-processing parameters (smoothing steps 23 or 28 instead of default 25) slices 263, 626, 628, 631, 690, 839, 859, 860 and 864 (in the substack with 1200 slices) to avoid whole-slice masks. The 3D reconstruction was performed in Blender 4.2 using the microscopy nodes (Github, oanegros/MicroscopyNodes).

## Data availability

All datasets are publicly available on scientific databases (EMPIAR, BBBC) or challenge platforms (Kaggle). The dataset of Fig. 3 is currently being uploaded to a public database.

## Code availability

All source code is released under BSD-3-Clause license and freely available on Github [26].

## Acknowledgements

We thank Noan Deschamps-Chevyreva for fruitful discussions and advice on data visualization. AKR and JDecelle were supported by the ANR EPHEMER and the ERC consolidator grant SymbiOCEAN (101088661). Data acquisition was possible thanks to AtlaSymbio. AtlaSymbio is funded by the Gordon and Betty Moore Foundation (grant ID GBMF11532, 10.37807/GBMF11532). MS and FJ were supported by AI4Life, and the work presented here was performed as part of the AI4Life Open Calls. AI4Life receives funding from the European Union’s Horizon Europe research and innovation programme under grant agreement number 101057970. Views and opinions expressed are however those of the author(s) only and do not necessarily reflect those of the European Union or the European Research Council Executive Agency. Neither the European Union nor the granting authority can be held responsible for them.

## Authors contributions

MS, FJ and JD developed the method; MS, VG and JD developed the code; AKR and JDecelle acquired data; MS, DDN, JB, and JD analysed the data; DDN, FJ and JD wrote the manuscript; FJ and JD supervised the project.

