## Supplementary figures for "FeatureForest: the power of foundation models, the usability of random forests"

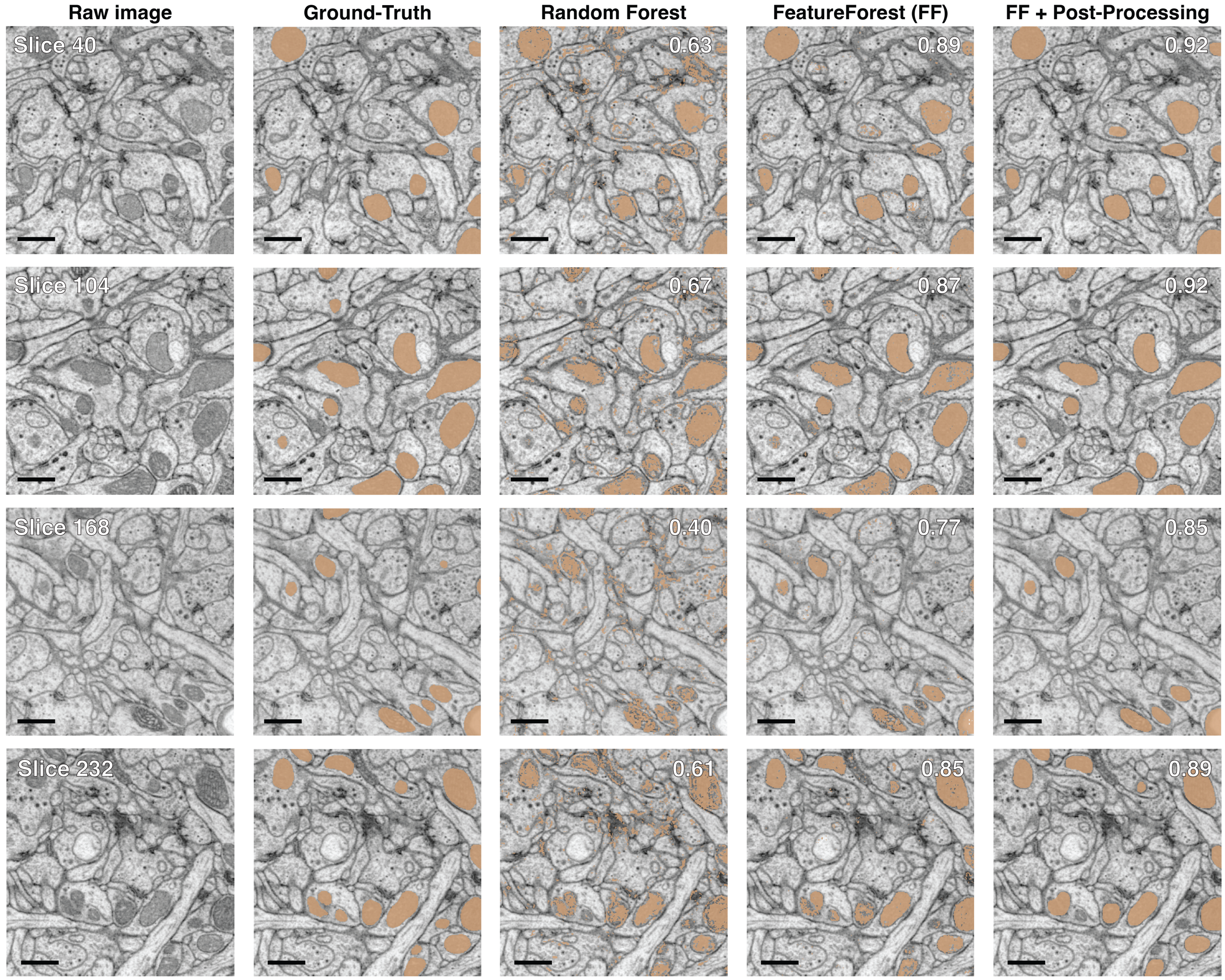


**Figure S1. Results on several slices of the FIB-SEM fly brain dataset.** FIB-SEM images of a fly brain, overlaid with mitochondria ground-truth mask, and with corresponding segmentations obtained with a random forest classifier, FeatureForest, and after post-processing the results from FeatureForest, from left to right respectively. Slice number is indicated on the top left corner of the raw image, and the Dice score with respect to the ground-truth for the specific slice and algorithm is indicated in the top right corner. Scale bars 500 nm.

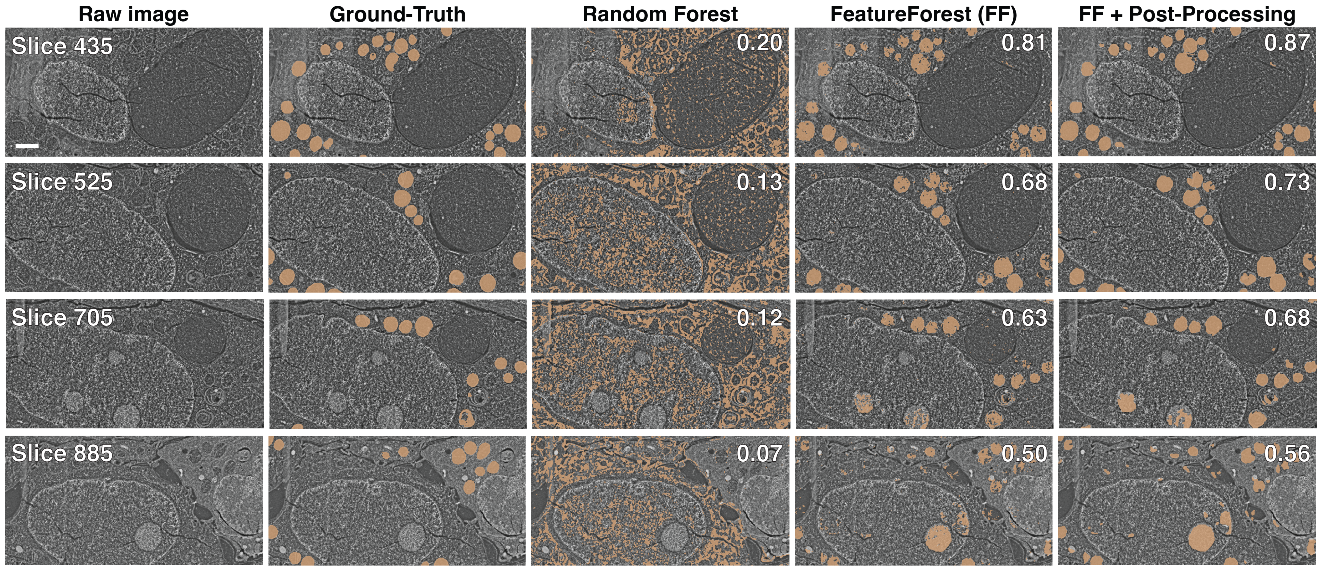


**Figure S2. Results on several slices of the FIB-SEM spheroid dataset.** FIB-SEM images of a human breast cancer spheroid, overlaid with mitochondria ground-truth mask, and with corresponding segmentations obtained with a random forest classifier, FeatureForest, and after post-processing the results from FeatureForest, from left to right respectively. Slice number is indicated on the top left corner of the raw image, and the Dice score with respect to the ground-truth for the specific slice and algorithm is indicated in the top right corner. Scale bars 1 µm.

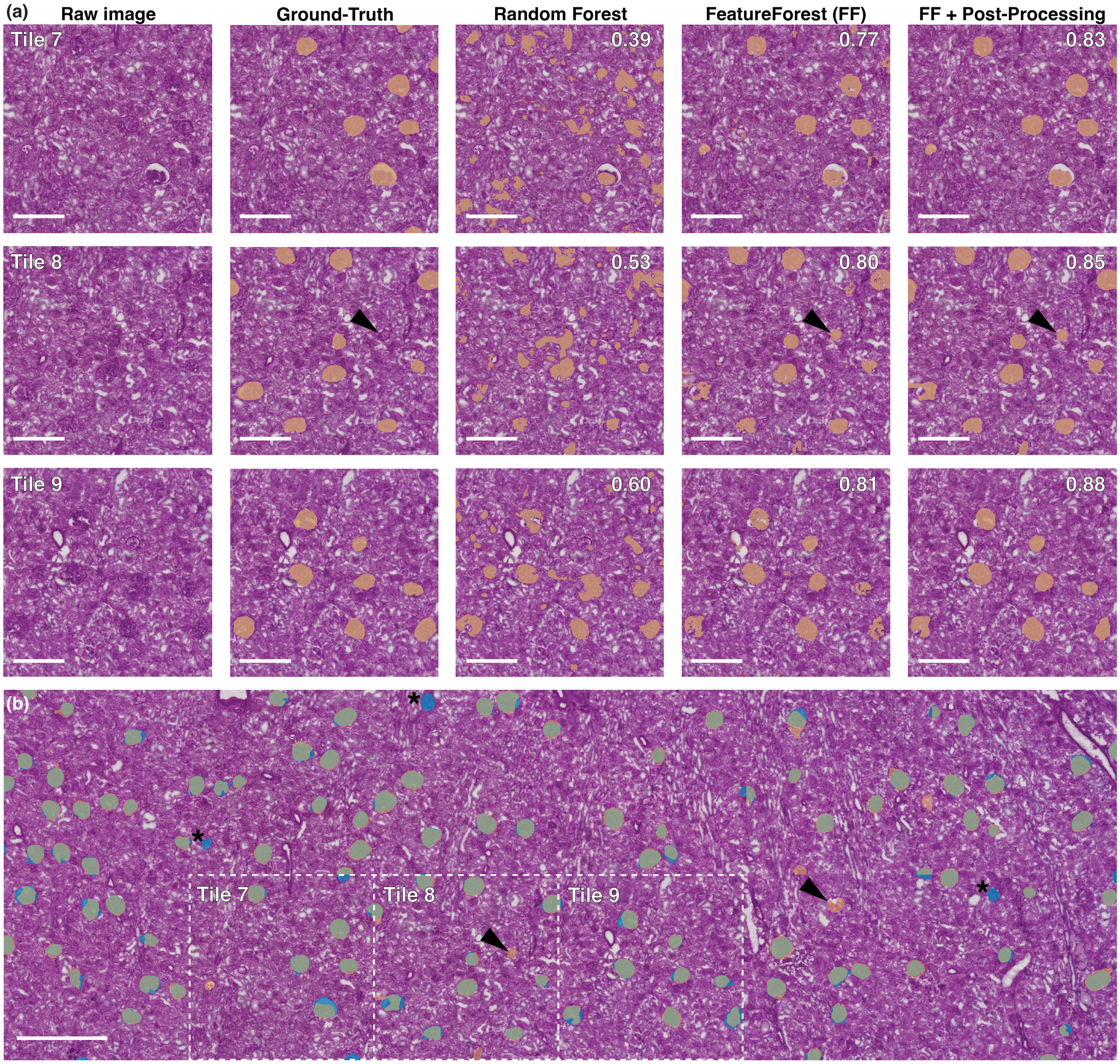


**Figure S3. Results on H&E stained human kidney dataset.** (a) H&E staining of a human kidney, overlaid with glomerulus ground-truth mask, and with corresponding segmentations obtained with a random forest classifier, FeatureForest, and after post-processing the results from FeatureForest, from left to right respectively. Arrows indicate a glomerulus instance segmented by FeatureForest and absent from the ground-truth. Tile number is indicated on the top left corner of the raw image, and the Dice score with respect to the ground-truth for the specific slice and algorithm is indicated in the top right corner. (b) Larger image from which the tiles are extracted with an overlay of both the ground-truth (blue) and FeatureForest + post-processing (orange). As in (a), arrows indicate an error in the ground-truth that is correctly segmented by FeatureForest. Asterisks label instances that are present in the ground-truth and absent in the FeatureForest segmentation. Tiles of (a) are delimited by the dotted rectangles. Scale bars 500 µm.

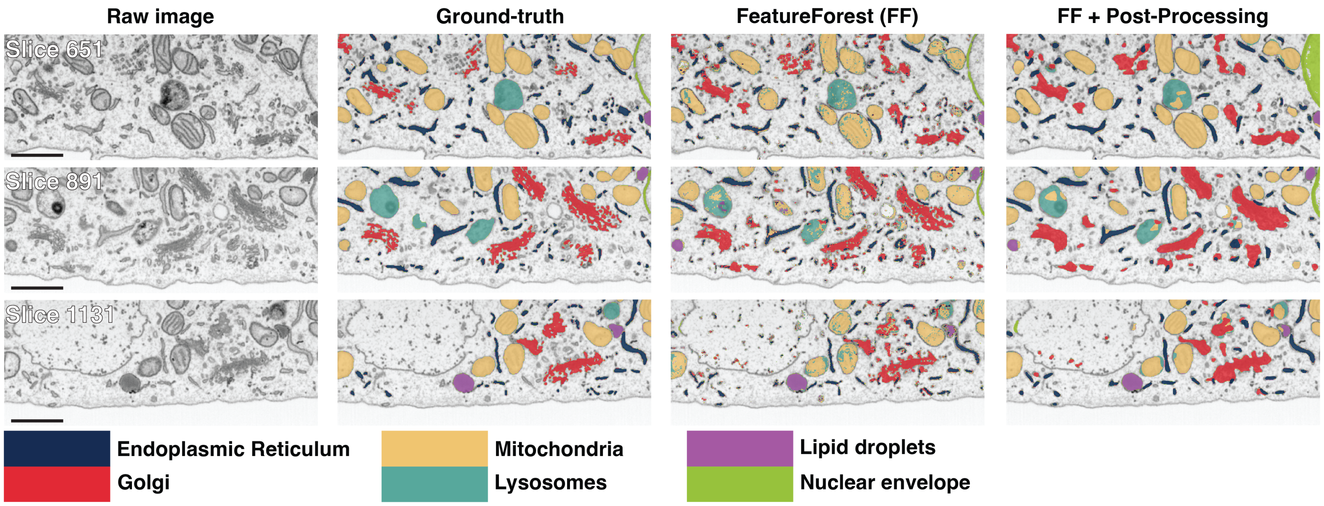


**Figure S4. Results on several slices of FIB-SEM images of a U2OS cell.** FIB-SEM images of a U2OS cell, overlaid with ground-truth for 6 different classes (see legend), and with corresponding segmentation obtained with FeatureForest, and after post-processing the results from FeatureForest, from left to right respectively. Slice number is indicated on the top left corner of the raw image. Scale bars 1 µm.

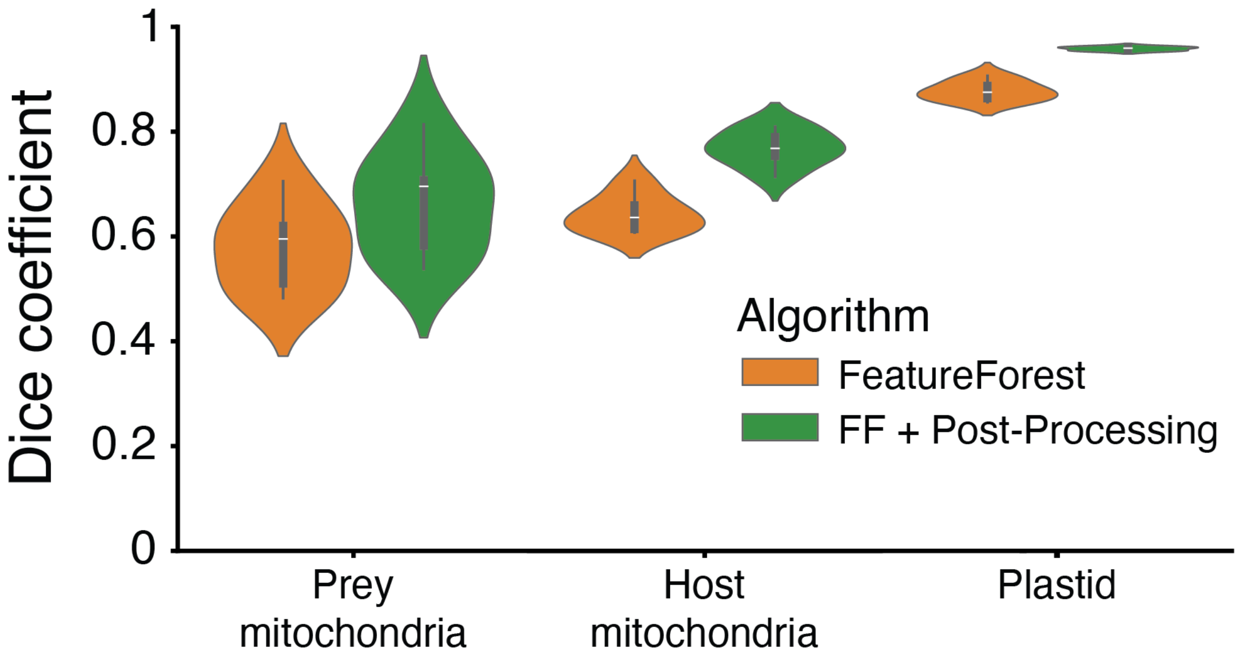


**Figure S5. Dice score over the test stack for the Dinoflagellate dataset.** Distribution of the Dice score over the test stack (7 slices) for FeatureForest and FeatureForest with post-processing for the three organelle classes.
